# Impact Of Learned Helplessness On Cognitive Performance And Resting-State Connectivity: An fMRI Study

**DOI:** 10.1101/2024.05.06.592810

**Authors:** Pierre Lechat, Eva Alonso-Ortiz, Jean-Samuel Bassetto

## Abstract

Learned helplessness (LH) is the phenomenon of resignation in the face of a problematic situation and is caused by the feeling of lacking control in a situation due to internal or external factors. This means that an individual does not seek to resolve the situation they are confronted with and consciously or unconsciously chooses to be passive. The Dorsal Raphe Nucleus (DRN) core could be at the root of the persistent action of the LH phenomenon and influences regions such as the striatum and amygdala. It is thought that the LH phenomenon could affect self-perception and it has been shown that the default mode network (DMN) is often associated with self-reflection during the resting state (RS) phase. It is therefore possible that LH influences both self-perception and the DMN. Based on previous studies, we investigated how LH affects participants. We used functional MRI (fMRI) to test this hypothesis. Participants were divided into two groups, subjected to solvable (control group), and solvable plus unsolvable (LH group) cognitive tasks. We also measured electrodermal signals, OCEAN (Openness, Conscientiousness, Extraversion, Agreeableness, Neuroticism) personality scores, variables related to the anagram resolution, and related these to LH. The study revealed significant differences in the RS contrast between the two groups, with the Posterior Cingulate Cortex (PCC) (an area of the DMN) being more connected to the DRN in the LH group than in the control group, and a portion of the Superior Temporal Gyrus being more connected to the PCC in the control group than in the LH group. These results suggest that LH may have a direct impact on the DMN and could lead to the start of long-term changes.

## 1.2 Introduction

Imagine an individual who, despite putting in consistent and dedicated efforts at their job, repeatedly faces unfavorable reviews and remains overlooked for promotions. These setbacks occur despite their qualifications and skills. Over time, this individual could believe that their actions cannot influence their professional situation, and they begin to experience a sense of powerlessness in the face of their circumstances. In this scenario, they may eventually choose to stop seeking improvements or opportunities for career growth, becoming passive in their approach to work-related challenges. This real-life example exemplifies the phenomenon of resignation and passivity often associated with learned helplessness (LH), where individuals perceive a lack of control over their circumstances due to internal or external factors, and as a result, they choose not to take action.

The LH phenomenon was initially discovered in animals when dogs no longer attempted to escape a stressful stimulus after learning that the stressful elements were inescapable (Maier et Seligman, 1976b). This phenomenon is also found in human beings. When an individual is faced with a perceived unattainable or unachievable goal, experiences a sense of powerlessness in the face of the consequences, and this situation becomes recurrent, they are likely to develop a state of LH. The study of LH in mice has shown that certain areas of the brain play a significant and measurable role in the phenomenon of LH. The Dorsal Raphe Nucleus (DRN) plays a clear role in the release of serotonin 5-HT, leading to the alteration of the functioning of areas such as the amygdala or the dorsal-periaqueductal gray (dPAG) (Maier & Seligman, 2016). The dorsomedial periaqueductal gray (dPAG) has been shown to play a role in the fight/flight/freeze system (Graeff et *al*., 1997) and the extended amygdala system (including the bed nucleus of the stria terminalis/stria terminalis) plays a role in the regulation of fear and anxiety (Graeff et *al*., 1997). Additionally, it has been demonstrated by Grahn et *al*. (1999) that the DRN is strongly activated during a stressful situation and remains active when the individual is exposed to a situation where no favorable outcome is perceptible. In other words, this region continues to be activated when the individual is in an uncontrollable situation but deactivates when the situation becomes controllable. Moreover, unlike the freeze phenomenon in the “fight/flight/freeze” system, which is an immediate reaction to a stressful stimulus, LH-induced passivity is generally associated with a slower reaction.

In their 2016 article, Maier et *al*. emphasize that the scientific community tends to believe that the activation of the DRN is both necessary (Will et *al*., 2004) and sufficient (Maier et *al*., 1995) to induce a state of passivity and anxiety in the subject. The necessity of DRN activation is demonstrated by blocking DRN activation, thereby inhibiting passivity and anxiety after an uncontrollable shock (Maier et *al*., 1993). Furthermore, Maier et *al*.’s 1995 article demonstrates that voluntary activation of the DRN using the neurotransmitter GABA induces behaviors in subjects similar to those observed in subjects suffering from LH. Thus, Maier et *al*.’s 2016 article indicates strong evidence of an equivalence between DRN activation and passivity/anxiety following an uncontrollable shock.

Several regions that have been identified as upstream of DRN activation can either inhibit or activate its effects. Notably, there are neural connections between the ventromedial prefrontal cortex (vmPFC) and the DRN that allow for its inhibition (Maier & Seligman, 2016). Additionally, the dorsomedial striatum (DMS) has connections with the vmPFC and acts in conjunction with the DMS to detect controllability. The DMS/vmPFC circuit is activated in escapable shocks but not in inescapable shocks (Amat et *al*., 2014), meaning that it detects controllability in escapable but not inescapable situations. This circuit also involves the substantia nigra (SN) and the mediodorsal thalamus (MD) (Baratta & Maier, 2019).

Translating the results of LH from mice to humans is challenging, especially since it involves measuring deep brain regions and may exceed ethical boundaries. To access these regions in mice, previous studies have employed various methods, such as direct electrophysiological measurements (Varela et *al.*, 2012), or in vivo microdialysis (Grahn et *al*., 1999). However, these methods are too invasive for use in humans. In addition, various harmful elements can be used to induce uncontrollable stress in humans, such as loud noises (Bollini et *al*., 2004; Henderson et *al*., 2012; Meine et al., 2020), electric shocks (Havranek et *al*., 2016), thermal stimulation (Bräscher et *al*., 2016), or insolvable tasks (Bauer et *al*., 2003). Regarding physical threats like uncontrollable noise disturbances, a recent study shows that participants subjected to LH are more exhausted and perform less well in subsequent trials where the physical threat is escapable (Meine et *al*., 2020).

It is important to introduce the default mode network (DMN) and why this network is important in our study. By definition, a neural network is identified when multiple regions of the brain exhibit a significantly correlated “Blood-oxygen-level-dependent imaging” (BOLD) signal over time. Initially, the DMN was discovered as the set of areas deactivating during a cognitive task (Shulman et *al*., 1997). Regarding its localization, the DMN is composed of several key nodes, including the posterior cingulate cortex (PCC), the vmPFC, the angular gyrus (AG), the dorsolateral prefrontal cortex (dlPFC), the inferior frontal gyrus (IFG), and other smaller areas (Anticevic et *al*., 2012). This neural network has garnered particular attention since dysfunctions within it have been identified in various psychological disorders such as attention deficit hyperactivity disorder, obsessive-compulsive disorder, post-traumatic stress disorder, and schizophrenia (Sha et *al*., 2019). In other words, connectivity analysis of the DMN has revealed notable differences between a healthy control group and a group affected by one of the mentioned disorders.

Numerous studies investigate the DMN by instructing subjects to engage in unfocused thinking during functional magnetic resonance imaging (fMRI) sessions, a sequence known as “resting-state” fMRI (rs-fMRI). Originally conceived to activate this network and distinguish brain functioning during a task from a resting period (Raichle et *al*., 2001), it has been demonstrated that this state persists not only when no specific task is assigned but also when the subject engages in self-centered (Molnar-Szakacs & Uddin, 2013) and/or emotional thoughts (Gusnard et *al*., 2001; Satpute & Lindquist, 2019). Indeed, even in the absence of a specific task, the participant may be reflecting on themselves during these moments.

Between 2011 and today, many other cognitive processes that activate the DMN have been identified, such as the ones mentioned below. This network is notably engaged during memory recall and the resurgence of autobiographical memories (Yang et *al*., 2013), mental simulation of future problem-solving (Gerlach et *al*., 2011; Pearson, 2019), moral judgments (Marín-Morales et *al*., 2022), and various other cognitive tasks involving self-projection. These cognitive processes ideally manifest when the mind is allowed to wander, particularly during a resting state.

This study replicates an experiment initially conducted in a classroom setting by Charisse Nixon (Paul, 2020). In this experiment, each student is given three anagrams to solve, and they are instructed to raise their hand when they find the answer. However, half of the class is given two unsolvable anagrams, while the other half is given only solvable. The third stage involves giving the same anagram to everyone. This classroom experiment reveals remarkable results cited by (Paul, 2020). The portion of the class that had to solve two unsolvable anagrams performs significantly worse than the other group when attempting the solvable anagram. Furthermore, students in this same group appear to exhibit the known characteristics of LH, namely passivity and resignation (e.g., showing signs of not thinking to find a solution, such as staring into space). Thus, it has been postulated that social comparison has a strong impact on LH. If the DRN acts in a similar way in both mice and humans, and if an unsolvable anagram has the same effect on the DRN as an uncontrollable shock, we hypothesized that the DRN may be activated following unsolvable anagrams and, thus, induce passivity in some individuals (less anagrams found, longer time of resolution). Moreover, we hypothesized that the DRN would deactivate when the anagram was found, since controllability was detected. This result should be particularly true in the control group, since many anagrams should be found. Finally, we have made the assumption that the maps resulting from the rs-fMRI analyses will be different between the two groups. It is possible that the DMN is altered following an LH experience, since LH could generate an alteration in self-perception.

To address these hypotheses, we designed an experimental protocol that used fMRI and was created accordingly to induce LH to participants from the LH group and not to the control group. This study explores the activation in the DRN while subjecting participants to solvable and unsolvable cognitive tasks. We also related physiological data such as electrodermal signals and the response to the OCEAN (Openness, Conscientiousness, Extraversion, Agreeableness, Neuroticism) scores to consequences of LH. Finally, we explored the differences in rs-fMRI between participants experiencing LH effects and those in the control group.

## 1.3 Methods

### 1.3.1 Data Acquisition

The 14 healthy participants (6 females and 8 males, M_age_ = 24 years, SD_age_ = 2 year) were recruited and randomly assigned to a control group (2 females, 5 males, M_age_ = 24 years, range 20-28) or LH group (4 females, 3 males, M_age_ = 24 years, range 22-25). All participants had normal or corrected-to-normal vision, normal hearing, were proficient in written and spoken French, were native French speakers and had no exclusion criteria for an MRI. The study was approved by a local Research Ethics Committee and all participants provided written consent before participating in the study. Derived data and codes for figures, preprocessing and analyses can be found at https://github.com/PierreLechat/article_codes_LH_fmri.git. Prior to the imaging experiment, participants were subject to the OCEAN test (Plaisant et *al*., 2010), i.e., a test that self classifies individuals according to five major personality traits: Openness to experience, Conscientiousness, Extroversion, Agreeableness, and Neuroticism.

Participants were scanned on a 3T scanner (Siemens Prisma-fit, Erlangen, Germany) equipped with a 64-channel phased-array head and neck coil. Two disposable MRI-compatible electrodes (https://www.biopac.com/product/disp-rt-dry-electrode-100pk/) were fixed on the inside of the right foot, approximately 8 cm apart, to measure electrodermal signals in participants using the BIOPAC MP160 measurement equipment and Acknowledge software (https://www.biopac.com/product/mp150-systems-with-ndt/) (the sampling frequency was set 100 Hz). Electrodermal data was preprocessed, and the phasic component was extracted using the default parameter of the module “eda_process” in the package Neurokit2 (Makowski et *al*., 2021) available in python. The imaging protocol consisted of a high-resolution T1-weighted (T1w) anatomical scan acquired using the MP2RAGE sequence (Marques et *al*., 2010) used for tissue segmentation and task-based and rs-fMRI acquisitions. Task-based fMRI was acquired as a series of 5 repetition blocks (with breaks between each block) using 2D multi-slice T2*-weighted (T2*w) echo planar imaging (EPI) acquisition (repetition time (TR) = 2890 ms, echo time (TE) = 23 ms, flip angle = 83.4°, matrix = 96 x 92, resolution = 2.1 mm x 2.1 mm, 40 slices, 2 mm slice thickness, bandwidth (BW) = 2004 Hz/pixel, partial Fourier phase = 6/8, total scan time = 317.9s, number of EPI for each block = 22, anterior-posterior (AP) phase encoding). Three EPI scans with the same parameters, but the opposite phase-encoding direction (posterior-anterior (PA)) were additionally acquired after the 5 blocks acquisition. Dual-echo gradient-echo fieldmap images were acquired to allow correction of the echo planar imaging distortions (TR = 650 ms, TE1 = 2.97 ms, TE2 = 5.43 ms).

During fMRI data acquisition, participants were asked to solve both solvable and unsolvable anagrams (LH group) or only solvable anagrams (control group).

Participants in the LH group had to complete five sets of three anagrams. Among these three anagrams, two were unsolvable (none in the control group), and only one was solvable (all three are solvable in the control group), which is the same as the third one in the control group. Figure 1 shows in (a) the construction of a block comprising the presentation of three anagrams for a duration of 20 seconds, then a white screen during which the participant’s anagram answers are requested, followed by the display of the other participants’ anagram results and finally the hint to signify passage to the next step. Part (b) of Figure 1 shows the acquisition of EPIs during the test, visually displaying the presses on the participant’s answer box.

**Figure 1.**
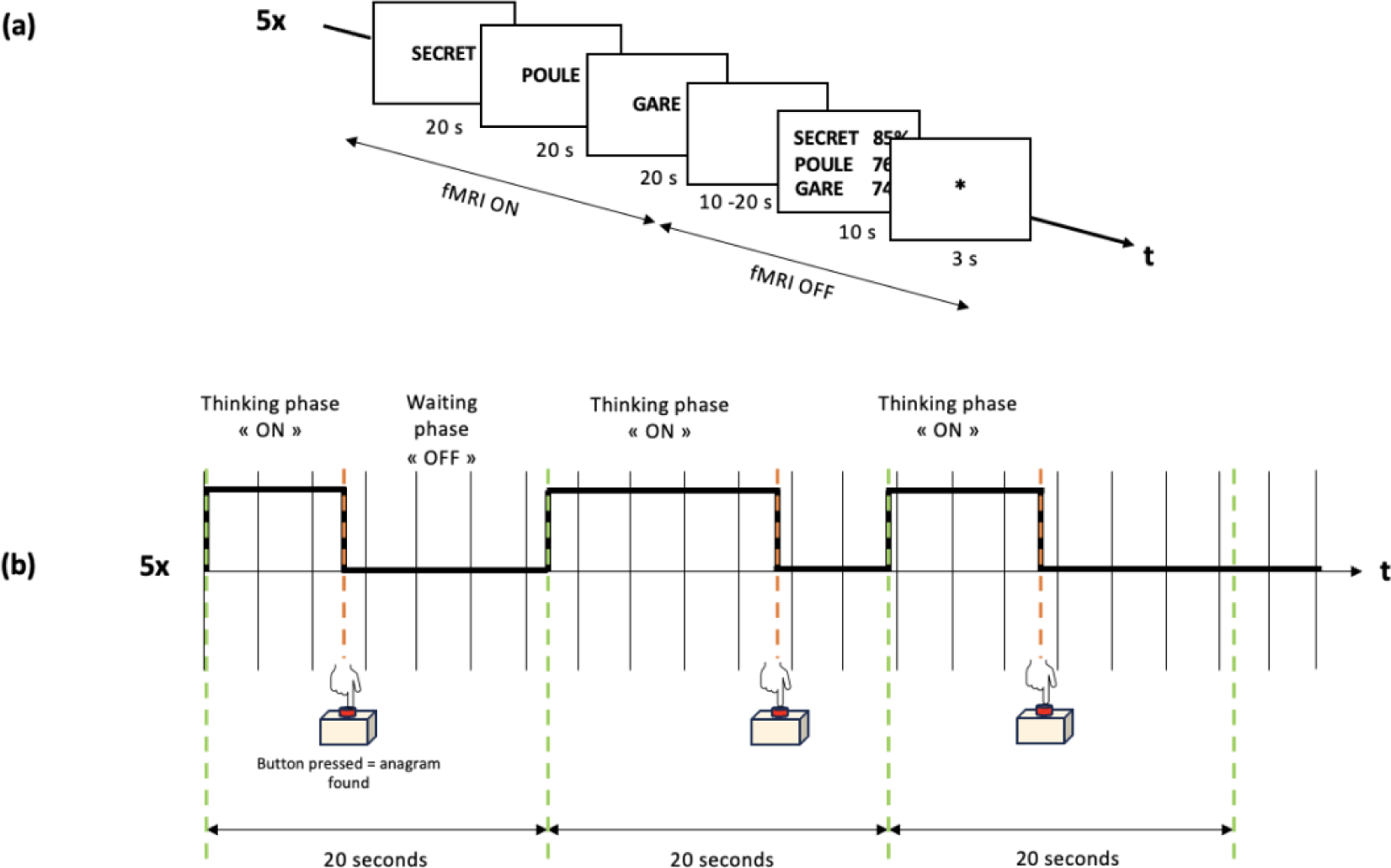
**(a)** fMRI scans are acquired during the resolution phase (i.e., during the presentation of the three anagrams, each displayed for 20 seconds). One task-based sequence block comprises 22 EPIs (i.e. 63.6s, to leave a margin of around 4s). Following this, a blank screen is presented, and participants are asked to provide their answers to the anagrams. Subsequently, the scores of other participants are displayed for 10s. Finally, a visual indicator “*” signifies that the sequence will restart in 3 seconds. **(b)** Illustrative example of the sequence in which the participant tried to find the solution to the anagrams. When they found the solution, they were asked to press the button they had. The vertical lines represent the TR times for acquired EPI volumes. The “thinking phase” represents the phase during which the participant tries to find an answer to the anagram and the response time is the duration of the “thinking phase”. The “waiting phase” represents the phase when the participant has already pressed the button, found the answer and is waiting for the 20s to elapse before moving on to the next part of the test. The terminology “resolution phase” will be used to describe one entire block of three anagrams (i.e, “thinking phase” and “waiting phase” combined).

Participants were asked to press a button on a response box once they had found the solution to a presented anagram. Pressing the button did not change the screen, and the participant had to wait 20s before proceeding to the next anagram. This 20-second time period was based on previous studies that examined anagram resolution in fMRI (Aziz-Zadeh et *al*., 2009), as well as the average anagram resolution time of 8 seconds that has been reported (Sinitsyn et *al*., 2020). Following the presentation of three anagrams, a blank screen would appear, and the responses were requested from participants (MRI data acquisition was paused during this period). Answers were given verbally to the experimenter via microphone. Then, a success rate for each of the previous three anagrams was displayed for 10 seconds, meant to reflect the success rate of previous participants. The purpose of the success rate presented after each sequence is to make the subject believe that other participants have already taken the test. In reality, the displayed success rates are fictitious. The goal was to convey to the subject that other participants have been successful in solving anagrams. It is assumed that this information will amplify the LH phenomenon in the LH group subjects (Fincham & Hokoda, 1987). Indeed, the choice of presenting fictitious success rates to the participants may imply that the anagrams have an answer, and thus induce the attribution of failure to internal rather than external causes. Internal, since it is the participants’ own “non-competence” that would induce failure at the anagrams. Moreover, it has already been shown that a social defeat can also more likely induce LH (Amat et al., 2010). Here we also verified participants’ responses for each anagram sequence since accidental button presses can easily occur. At the end of the 10 seconds time, a small star was shown to indicate that a new anagram would be presented in 3 seconds. This sequence of steps was repeated 5 times.

The rs-fMRI imaging protocol consisted of 2D T2*w EPI scans (TR = 2950 ms, TE = 29 ms, flip angle = 83.7°, matrix = 88 x 88, BW = 1536 Hz/pixel, partial Fourier phase = ⅞, resolution = 2.5 mm x 2.5 mm, 39 slices of 3.5 mm thickness, total scan time = 295 s, AP phase encoding), followed by 3 additional EPI scans with the same parameters but PA phase-encoding. Followed again by a dual-echo gradient-echo field mapping sequence (TR = 650 ms, TE1 = 2.97 ms, TE2 = 5.43 ms). During the rs-fMRI acquisition, participants were presented with a gray screen and instructed to let their minds wander without falling asleep.

After the fMRI and rs-fMRI scanning session, LH participants did not feel “competent” in solving anagrams, which is entirely normal since no answers could be found for some of them. Therefore, participants were told that they had just participated in a LH experiment, and it is normal that they felt frustration during and after attempting to solve the different anagrams. The purpose of this post-experience interview was to defuse any frustration and potential continuation of LH caused by the test they had just undergone. A set of positive phrases and vocabulary were provided to the participants for this purpose, such as: “It is entirely normal that you did not succeed since the anagrams were partly impossible.”.

### 1.3.2 fMRI Data Pre-Processing

MRI data were preprocessed using the FMRIB Software Library (FSL) v6.0, (Jenkinson et *al*., 2012). Steps included head motion correction using the MCFLIRT tool (Jenkinson et al., 2002), non-brain structure removal utilizing BET (S. M. Smith, 2002), susceptibility distortion correction based on AP/PA acquisitions (FSL’s topup module (Andersson et *al*., 2003)), correction of B0 distortions using B0 map (FSL’s FUGUE module (S. M. Smith et al., 2004)), and spatial smoothing using Gaussian kernel with 8 mm full width at half maximum (FWHM). Then, the fMRI scans were aligned to the subject’s high-resolution T1w image (i.e., MP2RAGE) using the boundary-based registration (Jenkinson & Smith, 2001). Finally, the T1-aligned fMRI images were non-linearly co-registered to the MNI152 template image (Andersson & Jenkinson, 2007).

### 1.3.3 fMRI Statistical Analysis

#### Task-based analysis

The statistical analysis of the task-based fMRI data was performed using the FMRI Expert Analysis Tool (FEAT) tool (Woolrich et *al*., 2004a), part of FSL, utilizing the general linear model (GLM). At the single-subject (i.e., first) level, EPI voxel time courses were correlated with the canonical Hemodynamic Response Function (HRF) convolved with the experimental event blocks (Figure 1b) resulting in 1 contrast of parameter estimate (COPE) map for each participant in each block for each group (LH and control group). A second level analysis for the control group was carried out using a mixed effect model (FLAME 1, (Woolrich et *al*., 2004b)), part of FSL, with the COPE image from all subjects from the control group. Z-statistic images were thresholded using clusters determined by z > 1.8 and a corrected cluster significance threshold of *p* = 0.01 was considered as significant. Then, a second-level analysis for the LH group was carried out using a mixed effect model, with the COPE image from the LH group as input. Z-statistic images were thresholded using clusters determined by z > 3.75 and a corrected cluster significance threshold of *p* = 0.1 was considered as significant. A group-level analysis was then carried out to compare the activation of the DRN in the “thinking phase” between the LH and the control group. Z-statistic images were thresholded using clusters determined by z > 2.3 and a corrected cluster significance threshold *p* < 0.01 was considered as significant.

#### Seed-based analysis for rs-fMRI

For the rs-fMRI data, we used the same preprocessing steps used for fMRI EPI data. We conducted a connectivity-based analysis using FEAT and the PCC, which was previously used as a seed of interest (Vatansever et *al*., 2017; Zhou et *al*., 2020). For this, the signal intensity in the seed of interest was averaged and a whole brain analysis was performed by comparing the intensity in the seed base with all other voxels. Data processing was made according to two levels. At the first level, EPI voxel time-courses were correlated with the average signal in the seed of interest. A second-level analysis at the group level was carried out using a mixed effect model. Z statistic images were thresholded using clusters determined by z > 1.8 and a corrected cluster significance threshold of *p* = 0.01 was considered as significant.

The various regions of interest mentioned in the introduction, such as the DRN, the caudate nucleus, the amygdala, the vmPFC and the PCC, were identified and segmented using the atlases cited below. The DRN region was segmented using the Levinson-Bari Limbic Brainstem Atlas, which includes the dorsal Raphe nucleus mask (Levinson et *al*., 2023). The caudate nucleus of the striatum was segmented using the WFU-Pick atlas (Version 3.3, Wake Forest University, School of Medicine, Winston-Salem, North Carolina; www.ansir.wfubmc.edu). The PCC has been segmented using the “Harvard-Oxford Subcortical Structural Atlas” available in FSL. The vmPFC was segmented using the “Harvard-Oxford Cortical Structural Atlas” (also available in FSL). These atlases have enabled us to identify the different activated clusters.

### 1.3.4 Statistical analysis for the other variables

Statistical analysis for variables related to the anagram test such as response time, score, number of instruction errors, OCEAN test variables, and number of peaks in the electrodermal signal was performed using the scipy “stats” module in Python (Virtanen et *al*., 2020). Instruction errors mean that the participant has not followed one of the rules presented before the session, i.e., pressing the answer box several times for a single anagram, pressing the answer box when no answer has been found, giving an answer in English or forgetting an answer that had been found.

These different methodological points allow us to verify in the following section our different hypotheses such as the relationship between OCEAN test variables and LH, between electrodermal signal and LH, between anagram resolution and LH and the observability of difference between resting state maps between the two groups.

## 1.4 Results

### 1.4.1 OCEAN test

The verification of an unbiased group in the OCEAN test is essential to judge the real effect of LH alone. Indeed, unbalanced groups in terms of OCEAN test variables could attenuate or accentuate the results that a balanced sample would allow. We therefore measured the intercorrelations between the OCEAN test variables of all subjects and how they were distributed between the control and LH groups. First, a Shapiro-Wilk test (S) revealed that the OCEAN test variables (S_Openness_(12) = 0.95, *p*_Openness_= 0.53, S_Conscientiousness_(12) = 0.94, *p*_Conscientiousness_ = 0.38, S_Extraversion_(12) = 0.95, *p*_Extraversion_ = 0.56, S_Agreeableness_(12) = 0.94, *p*_Agreeableness_ = 0.36, S_Neuroticism_(14) = 0.93, *p*_Neuroticism_ = 0.27) and the response time (SRT(49) = 0.84, *p*response time < 2.10^-5^) (where “Sx(y) is the statistic, “x” is the OCEAN test variable, and “y” is the degree of freedom) were not significantly normally distributed so we decided to perform a Spearman (non-parametric statistical test) coefficient correlation calculation (r). Several assessment axes of the OCEAN test revealed high significant correlations (Figure 2) (Spearman correlation coefficients: r_Openness/Agreeableness_ (12) = 0.77, *p*_Openness/Agreeableness_ < 0.0015, r_Agreeableness/Extraversion_(12) = 0.78, *p*_Agreeableness/Extraversion_ < 0.001). Conversely, there appeared to be a medium negative correlation between Extraversion and Neuroticism, and this was found to be statistically significant (r_Extraversion/Neuroticism_(12) = −0.55, *p*_Extraversion/Neuroticism_ < 0.05). Figure 2 shows the matrix of intercorrelations between variables in the OCEAN test, using a colored gradient from bright for high positive correlation to dark for strong negative correlation.

**Figure 2.**
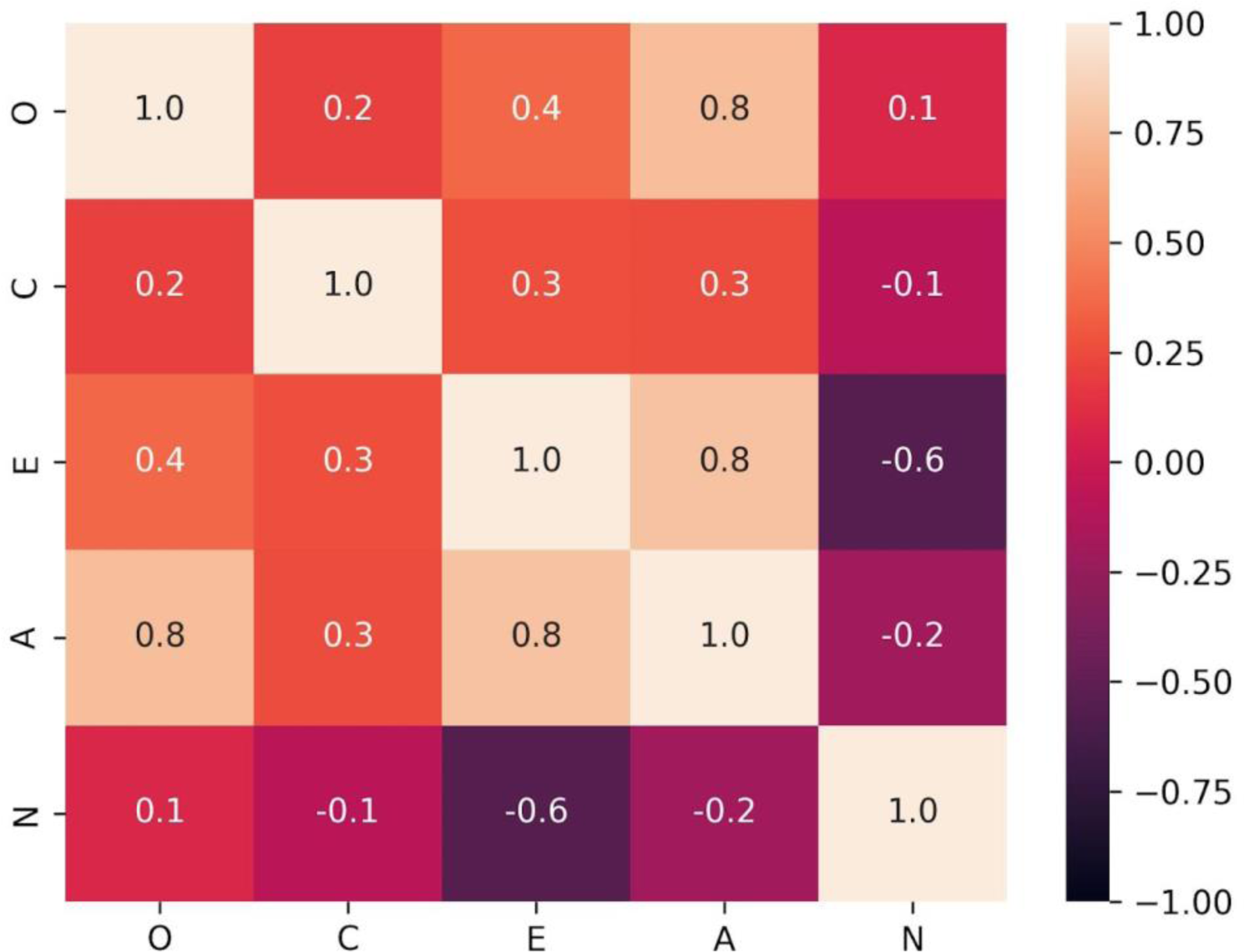
Spearman correlation matrix showing the correlation coefficient between the various variables of the OCEAN personality test, including Openness (O), Conscientiousness (C), Extraversion (E), Agreeableness (A), and Neuroticism (N), for all subjects.

#### Spearman correlation coefficient matrix of the OCEAN test variables for all subjects

The Kolmogorov-Smirnov (ks) test revealed that the control and LH groups were not all significantly well balanced according to OCEAN scores (ks_Openness_(12) = 0.43, *p*_Openness_ = 0.58, ks_Conscientiousness_(12) = 0.14, *p*_Conscientiousness_ > 0.99, ks_Extraversion_(12) = 0.29, *p*_Extraversion_ > 0.95, ks_Agreeableness_(12) = 0.43, *p*_Agreeableness_ = 0.58, ks_Neuroticism_(12) = 0.29, *p*_Neuroticism_ > 0.95). These results will be considered in the analysis of the consequence of LH on the variables relative to the anagram resolution (response time, proportion of correct answers).

### 1.4.2 Effect of LH on solvable anagram resolution

We measured the differences between the control and LH groups for several variables related to anagram solving (i.e., proportion of correct answers, response time and number of instruction errors). It’s important to note that these values are always derived from the same anagrams solved between the two groups.

Concerning anagram resolution scores, the average (µ) anagram resolution scores were found to be: µ_LH_ = 3.43, SD_LH_ = 1.27, µ_control group_ = 3.71, SD_control group_ = 0.95, for LH and control groups respectively (SD is the standard deviation). A **χ**^2^ test between the two samples revealed non-significant differences (**χ**^2^(12) = 0.28, *p* > 0.58). Furthermore, the average number of instruction errors per participant was higher in the LH group, although this was found to be nonsignificant while conducting a Mann-Whitney test (U) (µ_control group_ = 1.86, µ_LH_ = 3.00, U(12) = 16.5, *p* = 0.32). Moreover, these results should be interpreted with caution. More than 10% of the instruction violations in the LH group resulted from the response given to the anagram “GERME,” which led to the response “MERGE” in English. This might have been an oversight by the participants who provided this response. The differences in instruction violations between the two groups are less significant when this error is not counted (µ_control group_ = 1.86, µ_LH_ = 2.43, U(12) = 20.5, *p* = 0.65).

Regarding response time, it was observed that the average supposed response time is significantly greater in the LH group than in the control group (µ_response time-control group_ = 8.99 s, µ_response time-LH_ = 13.15, ks(12) = 0.39, *p* < 0.006). One might question whether the 20-second thinking time, when no response is given by the participant, artificially increases the difference between the two groups. Therefore, it is interesting to examine the time difference between the two groups when a response is given. It turns out that the average response time for correct answers only is still higher in the group subjected to LH (µ_response time-solution found-control group_ = 5.17 s, µ_response time-solution found-LH_ = 9.88 s, ks(12) =0.51, *p* < 0.0025).

Finally, it is also noted that the number of self-deprecating comments is higher in the LH group than in the control group. We recorded some comments and reactions from participants subjected to LH:

- “Response ‘DEBILE’ given to the anagram ‘DECIBEL.’”
- “Laughter during the session since no answers were found.”
- Explanations provided such as: ‘I was almost there every time,’ ‘I’m not used to thinking anymore,’ ‘I should have practiced before coming.’.

### 1.4.3 Relationship between anagram resolution and OCEAN test

Here, we tested the relationships between variables related to participants’ responses during anagram resolution (response time, anagram scores, number of instruction errors) and their scores on the OCEAN test. In each of the cases below, we tested the hypothesis that the scores for Openness, Conscientiousness, Extraversion, and agreeability are not related to different response variables, and the hypothesis that the score for Neuroticism is inversely related to the response variables.

Regarding response time, we did not find a significant correlation with scores for Openness, Conscientiousness, Extraversion and agreability (r_response time/Openness_ = −0.11, *p*_response time/Openness_ = 0.38, r_response time/Conscientiousness_ = 0.06, *p*_response time/Conscientiousness_ = 0.63, r_response time/Extraversion_ = −0.06, *p*_response time/Extraversion_ = 0.64, r_response time/Agreeableness_ = - 0.13, *p*_response time/Agreeableness_ = 0.29). However, it appears that Neuroticism is weakly correlated with thinking time (r_response time/Neuroticism_ = 0.24, *p*_response time/Neuroticism_ < 0.05). Figure 3 shows the linears regression between response time and the 5 variables of the OCEAN test, with the control group data in green and the LH group data in red.

**Figure 3.**
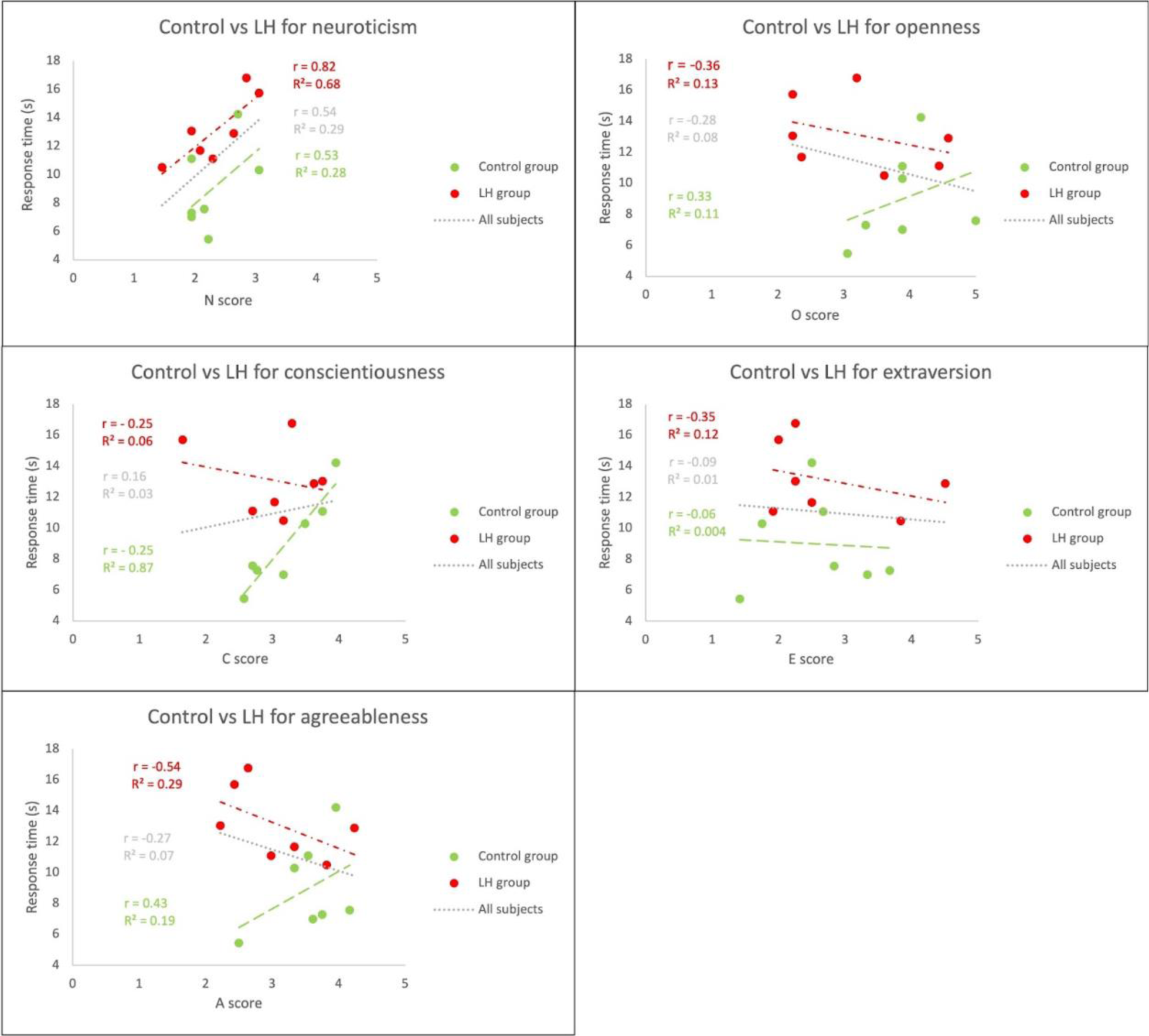
Relationship between the anagram response time and the various variables of the OCEAN personality test, including Openness (O), Conscientiousness (C), Extraversion (E), Agreeableness (A), and Neuroticism (N) separated between the groups. Control group subjects are shown in green, LH group subjects are shown in red, and all subjects grouped together are shown in gray.

Moreover, when considering only the binary value of the participant’s anagram response (i.e., 1 when a correct answer was found and 0 otherwise), there is a negative and significant correlation between a correct response and the Neuroticism score on the OCEAN test (r_response/Neuroticism_ = −0.29, *p*_response/Neuroticism_ < 0.013). Using this same method, no significant correlation is found with the other OCEAN test scores (r_response/Openness_ = 0.05, *p*_response/Openness_ = 0.69, r_response/Conscientiousness_ = −0.11, *p*_response/Conscientiousness_ = 0.37, r_response/Extraversion_ = 0.06, *p*_response/Extraversion_ = 0.64, r_response/Agreeableness_ = 0.08, *p*_response/Agreeableness_ = 0.52).

Concerning the number of instruction errors, we did not find significative correlation between the number of instruction error and OCEAN score as suggested by the following p-values (r_Openness/instruction errors_(12) = −0.25, *p*_Openness/instruction errors_ = 0.39, r_Conscientiousness/instruction errors_(12) = −0.14, *p*_Conscientiousness/instruction errors_ = 0.63, r_Extraversion/instruction errors_(12) = −0.39, *p*_Extraversion/instruction errors_ = 0.17, r_Agreeableness/instruction errors_(12) = −0.25, *p*_Agreeableness/instruction errors_ = 0.39, r_Neuroticism/instruction errors_(12) = 0.31, *p*_Neuroticism/instruction errors_ = 0.28).

### 1.4.4 Impact of LH on electrodermal activity changes

The focus here was on the differences in electrodermal signals during the resolution phases between participants in the control and LH groups and particularly on the number of peak differences. The LH group showed more peaks than the control group (N_peaks/LH/resolution phase_ = 250, N_peaks/control group/resolution phase_ = 190, where N is the number of peaks in the group during all the resolution phase). Moreover, a significant difference between the distribution was observed between LH group and control group in the number of peaks in participants’ electrodermal signals in the global session of task (ks(12) = 0.71, *p <* 0.05). This means that participants in the LH group tend to have more peaks in their electrodermal signal when performing the tests than the control group.

### 1.4.5 Activation of DRN and the anterior caudate nucleus in the control group during solved tasks

We are interested here in testing the hypothesis that the DRN activates during a stressful stimulus and deactivates when the controllability of the stimulus is detected. The activation pattern of the BOLD signal in the DRN and in an anterior part of the left caudate nucleus showed a strong correlation with the task “thinking phase” when doing a mixed effect analyses between the participants (z_max_ = 2.66, z_threshold_ > 1.8, *p*_cluster correction_ = 3.41 x 10^-2^, n_voxels_ = 14, and z_max_ = 2.96, z_threshold_ > 1.8, *p*_cluster correction_ = 2.33 x 10^-9^, n-voxels = 43, respectively). Figure 4 shows clusters with z above 1.8 superimposed on the MNI-152 template where structures such as the DRN and an anterior part of the left caudate nucleus appear to be activated during the “thinking phase” event.

**Figure 4.**
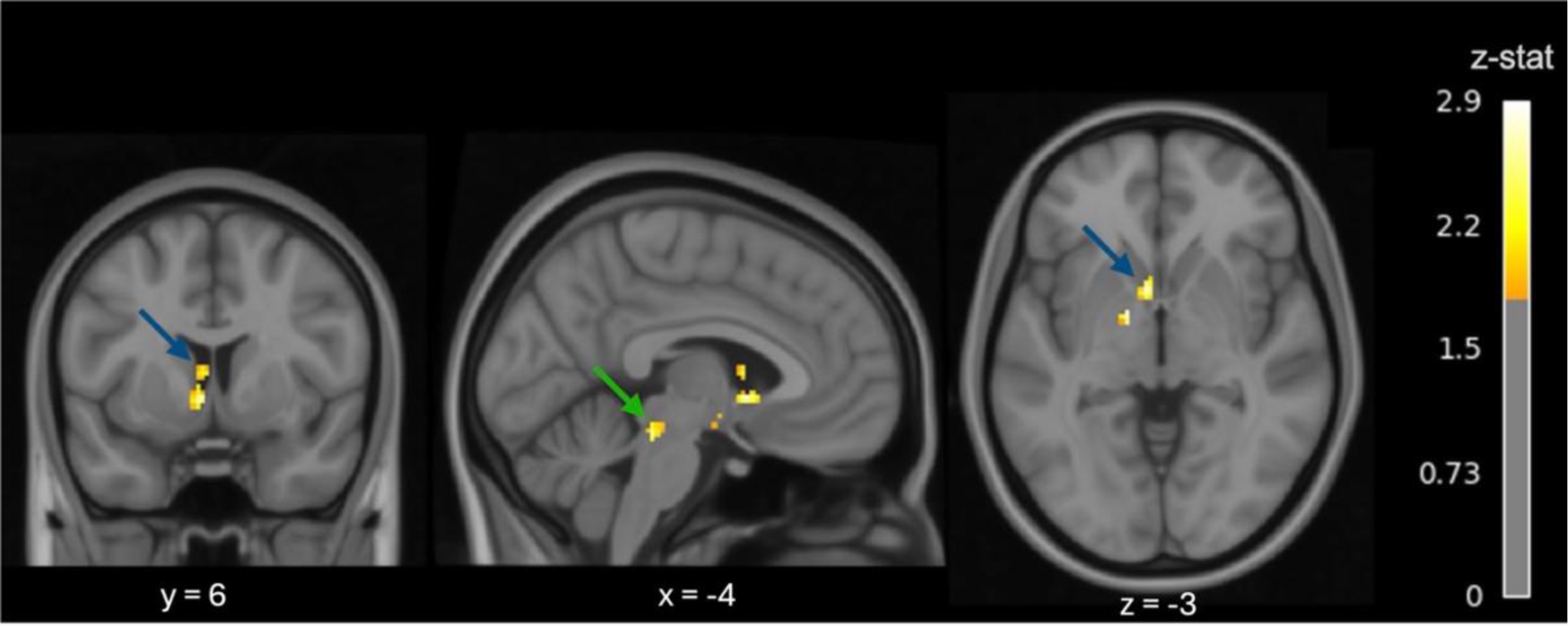
Z-statistic (thresholded at z < 1.8 and cluster corrected at p < 0.01) map for control group participants obtained from higher-level analysis carried out using a mixed effect model, superimposed on the MNI-152 template. A high z-statistic indicates a high correlation between voxel intensity over time and the event “thinking phase”. A high z-statistic is observed for voxels in the DRN and the caudate striatum. The DRN and the caudate striatum are indicated by the green and blue arrows, respectively.

### 1.4.6 Stronger DRN activation in the LH group during tasks

The focus here is on DRN activation in the LH group during the “thinking phase”. The task-based analysis based on the “thinking phase” event shows in Figure 5 that many areas had high z-scores, and the 6th most significant area was located in the DRN region (z_max_ = 4.69, z_threshold_ > 3.75, *p*_cluster correction_ = 0.0906, n_voxels_ = 19) meaning that there is a strong correlation between the event and the voxel time course. Figure 5 shows voxels declared activated (voxels in yellow to white) at z > 3.75 and a cluster correction at p > 0.1 signifying a strong correlation between voxel time course and the “thinking phase” event.

**Figure 5.**
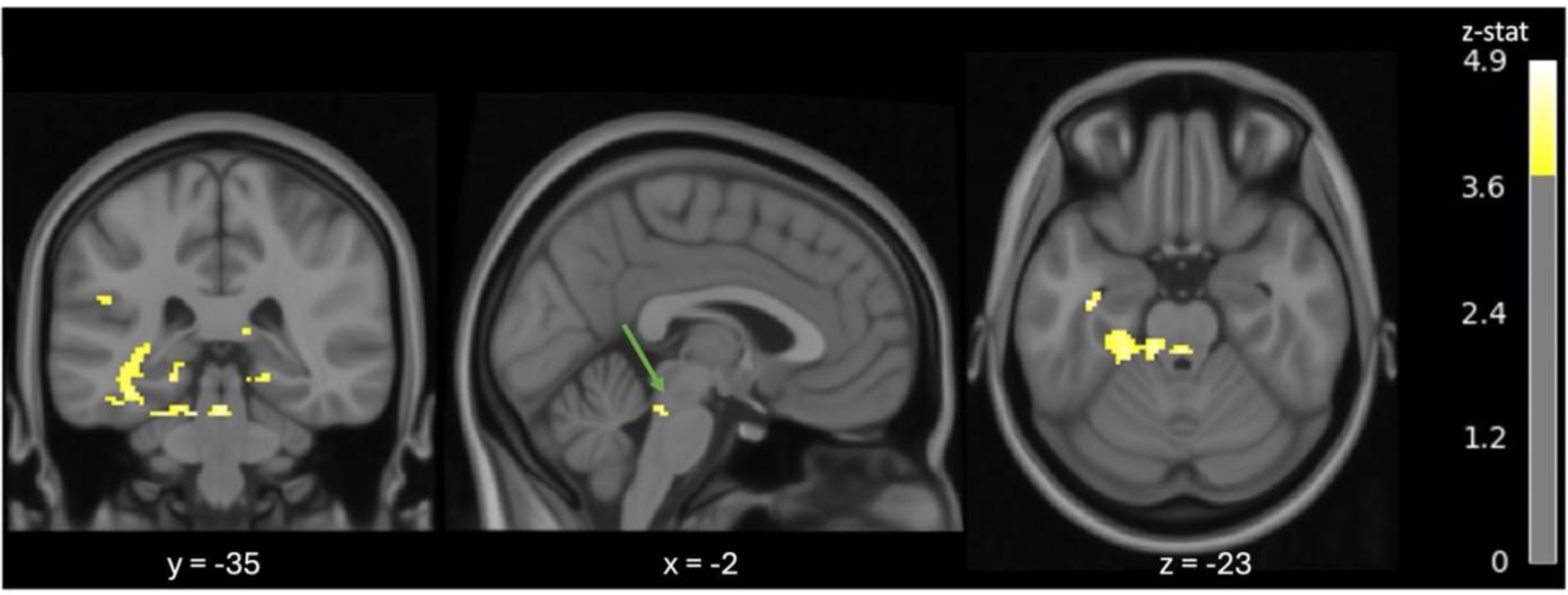
Z-statistic (thresholded at z < 3.75 and cluster corrected at p < 0.1) map for LH participants obtained from second-level analysis carried out using a mixed effect model, superimposed on the MNI-152 template. A high z-statistic indicates a high correlation between voxel intensity over time and the event “thinking phase”. One cluster appears to be in the DRN region which is indicated by the green arrow.

Subsequently, we compared the difference in the correlation between DRN activation for the LH group and the control group. The results show a greater positive correlation between the voxel time course and the event “thinking phase” in a portion of the DRN when contrasting the LH group by the control group as shown in Figure 6 (z_max_ = 3, z_threshold_ > 2.3, *p*_cluster correction_ = 6.08 × 10^-4^, n_voxels_= 10) meaning that voxels are more correlated to the event in the LH group than in the control group.

**Figure 6.**
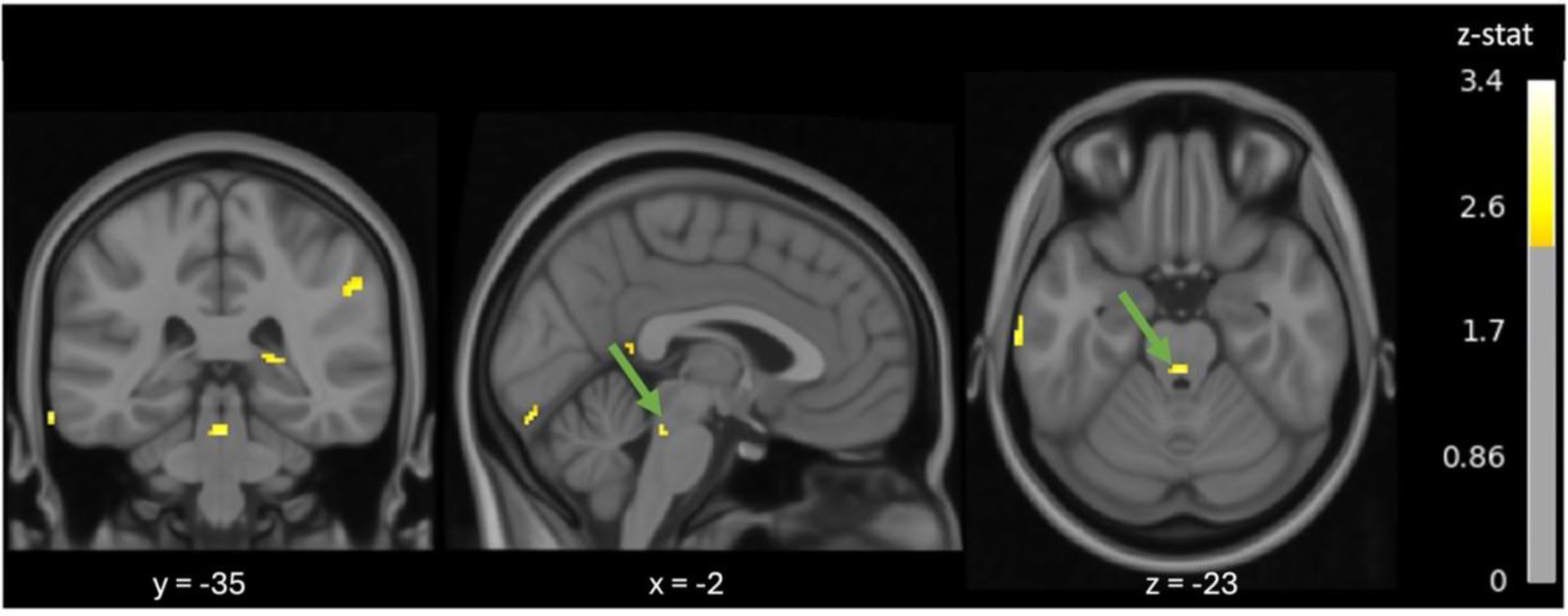
Z-statistic (thresholded at 2.3) map for LH and control group participants obtained from second-level analysis carried out using a mixed effect model and when contrasting the LH group by the control group, superimposed on the MNI-152 template. A high z-statistic indicates a large difference between LH and control group for the correlation between voxel intensity over time and the event “thinking phase”. One cluster appears to be in the DRN region, as indicated by the green arrow.

### 1.4.7 rs-fMRI differences between the two groups

We are interested here in exploratory research to test the hypothesis that RS may be different between the control group not subjected to LH and the LH group that has been subjected to LH and to explore the region where difference appeared. This analysis revealed differences in connectivity patterns in various brain regions between the control group and the group subjected to LH. In the DRN, we observed that the mean z-statistic was higher in the LH group (µ_z-score_ = 2.35, SD = 0.60) than in the control group (µ_z-score_ = 1.49, SD = 0.59). A Student’s t-test was conducted on this difference, resulting in a highly significant contrast (t(147) = 12.54, *p* < 1.10^-28^).

Moreover, in an extensive exploratory analysis of the whole brain, by contrasting the LH group with the control group and applying a cluster correction (p_cluster_ < 0.1), we found that regions exhibiting stronger connectivity with the PCC (i.e, high correlation between the mean average signal within the PCC with others voxels in the brain) in the LH group, compared to the control group (as highlighted by the voxels in red and yellow in Figure 7) were located within the DRN (indicated by the green arrow). Conversely, in the case of the control group, the posterior Superior Temporal Gyrus showed greater connectivity with the PCC (indicated by the blue arrow), compared to the LH group as highlighted by the blue voxels in Figure 7). Thus, Figure 7 shows the RS of the LH group contrasted by the control group, the resulting map of RS analysis being based on the strong correlation between the participants’ time course voxels with their temporal signal (spatial average) from the PCC. The statistical map is thresholded at z > 1.8 and z < −1.8 and a cluster correction is applied at p < 0.1.

**Figure 7.**
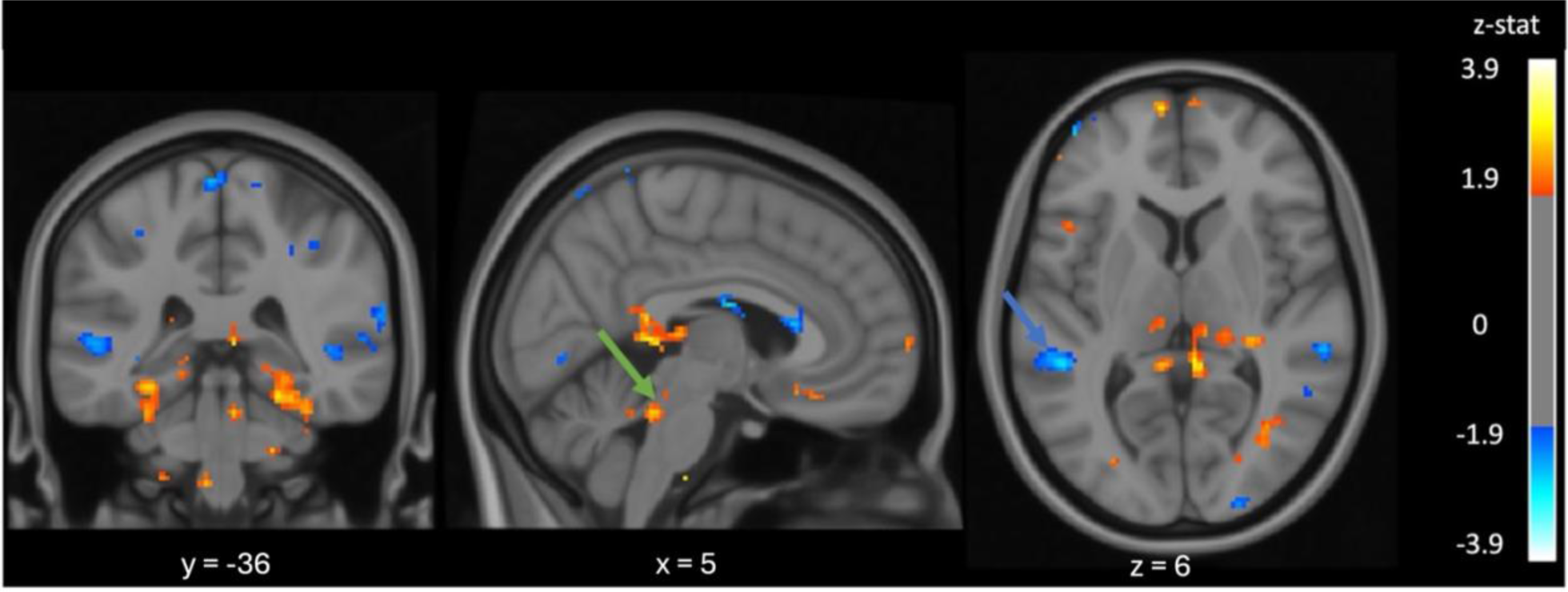
Z-statistic (thresholded at +/-1.8 and cluster corrected at p < 0.1) map for LH and control group participants obtained from higher-level analysis carried out using a mixed effect model, superimposed on the MNI-152 template. A high z-statistic (red and yellow voxels) indicates a high correlation between voxel signals from LH and the PCC seed signal and a high negative z-statistic (blue voxels) indicates a high correlation between voxel signals from control group and the PCC seed signal. One cluster appears to be in the DRN region in the LH group which is indicated by the green arrow and one cluster appears to be in the posterior superior temporal gyrus which is indicated by the blue arrow.

## 1.5 Discussion

The aim of this research was to study the effects of LH using a novel experimental setup and metrics: On one hand, we sought to understand the impact of personality traits on the effects of learned helplessness in a cognitive task and, on the other, to demonstrate the effect of LH on physiological phenomena. In addition, the aim of this study was to show that it was possible to measure global activation of the DRN during thinking phases. Finally, the main objective of this research was to show the lasting negative effect of LH over time and thus to demonstrate that, like trauma, changes in connectivity maps could be observed in the LH group versus the control group.

An important point to note, which limits our findings, is that the control group and the LH group are not properly balanced in terms of participants’ gender. Specifically, the control group consists of 5 men and 2 women, while the LH group comprises 4 women and 3 men. This disparity in participants’ gender may either attenuate or accentuate certain observed phenomena. Indeed, the RS differs between men and women (Filippi et al., 2013), potentially resulting in differences between the LH and control groups beyond the experience of LH. To better target the effects of IA, a later study could take care to have groups of participants made up of the same proportions in terms of gender.

Another important point for future studies using unsolvable anagrams in cognitive testing will be to check that the words given have no possible answer in other languages in which the participants might speak. In fact, one of the words given in French, “germe”, attracted responses in English with the word “merge”. This deviation from the initial protocol was taken into account in the statistical analysis, so that this case was not taken into account.

Regarding the results of the OCEAN test for the selected participants, it appears that certain measures from the test (Openness and Agreeableness, Agreeableness and Extraversion) are significantly positively correlated in the selected sample and Neuroticism is negatively correlated to Extraversion. A meta-analysis from 2010 combining the results of 212 OCEAN studies shows that variables among this test are correlated (Van Der Linden et *al*., 2010). Moreover, the study from Van Der Linden et al. shows that Neuroticism is negatively correlated to the other OCEAN variables when the other variables are positively correlated to the others. Our study has not found a significant negative correlation between Neuroticism and the other OCEAN variables (other than Extraversion) and not all OCEAN variables (other than Neuroticism) were intercorrelated. It’s possible that our sample is biased compared to one that would better characterize the overall human population (participants are young and mostly students). Increasing the number of older participants in diverse life situations may help reduce bias in the sample.

In the selected sample, the results of the OCEAN scores seem to be related to the variables concerning anagram resolution. Specifically, Neuroticism is significantly and negatively correlated with response time and the number of correct answers. This result confirms the finding of the study of Beckmann and al. in 2013 showing that a high score for Neuroticism is negatively related with performance (Beckmann et al., 2013). However, in our study data seems to only show a linear fit between Neuroticism and performance unlike the study of Beckmann *et al*. which found a quadratic relation between Neuroticism and task performance. A sample with a lower Neuroticism score can help to see whether a Neuroticism score below a certain value tends to decrease performance on the task, as in the study by Beckmann et *al*.. It is therefore possible that the effects of LH are accentuated by Neuroticism in individuals because people with higher neurotic scores might perceive the source of their failure as being more easily the result of their own incompetence. In consequence, they could be more passive after the first failure during the test and found the response slower. In addition, other personality characteristics seem to go in the direction of attenuating the harmful effects of LH. It’s possible that an agreeable person will more readily accept they are not perceived as performing well. Similarly, an open-minded person is more likely to accept their own failures and bounce back more easily in the future. It would therefore be interesting, by increasing the number of participants, to see whether Agreeableness and Extraversion have positive effects on the deleterious effects of LH. A simpler protocol experiment (without fMRI acquisition) could allow to have a larger sample.

Concerning the impact of LH on resolution variables, there is a significant increase in response time in the LH group. This suggests that our study succeeded in impacting participants in the LH group. There are also several self-deprecating responses and attitudes in the LH group. These results highlight the real and negative effect of LH during the task execution period, as suggested by previous studies (Meine et *al*., 2020; Valås, 2001). Moreover, it confirms that a simple task such as the resolution of unsolvable anagrams can lead to observable effects of LH.

In the same vein, there is a significant difference between the distribution in the number of sudden variations in the electrodermal signal of LH group and control group. Therefore, for the LH group, it’s possible that during phases where participants are exploring the nature of controllability, LH participants find it more difficult to obtain a response and thus experience more emotional variations.

fMRI revealed that a part of the DRN was more activated during the thinking phase of participants in the control group, which is in direct agreement with the article by Grahn et *al*. (1999). Indeed, since participants in the control group are more likely to find a response and the DRN is supposed to deactivate when controllability is detected, this justifies the fact that we do indeed obtain a good correlation between the event and the temporal signal in this region. Moreover, the prefrontal cortex is not declared activated during the resolution phase. It’s possible that the pre-frontal cortex doesn’t deactivate after the button is pressed and the anagram is solved. It could be that the participant checks his answer several times, and thus continues to think about the problem, without the DRN being activated, since controllability has been detected. Thus, the pre-frontal cortex would not be declared activated, since the “thinking phase” event would not characterize the activation/deactivation of the pre-frontal cortex. Nevertheless, this result needs to be looked at carefully. In fact, even if the cluster corrected threshold of p < 0.01 is a commonly used value (Chen et *al*., 2018; Woo et *al*., 2014; Yeung, 2018), the z statistic threshold of 1.8 is not and is more commonly set at 2.3 or even 3.1 and more (Cai et *al*., 2023; Criss et *al*., 2021). In addition, one of the limitations in our study was the low temporal continuity of participants’ data. Indeed, having to request responses from participants after every three anagrams proved to be impractical during data analysis. However, it was observed that, in the context of unsolvable anagrams, breaks are necessary to verify responses. It might be considered in another type of unsolvable cognitive test to conduct a single acquisition sequence, but one that is longer. Furthermore, in the LH group, it was observed that some of the voxels in the DRN are declared activated, meaning that we found a correlation between these voxels’ time course and the “thinking phase” event. However, the cluster correction threshold set at p < 0.1 is not a commonly used value. Indeed, this result needs to be looked at carefully. There are multiple regions that appear in the map that are not known, to the best of our knowledge, to be related to LH or to a stressful stimulus. The main limitation of our protocol in this case is that the “thinking phase” event may poorly predict DRN activation. In fact, it was assumed in the beginning that at the start of each new acquisition sequence, the DRN would be deactivated and reactivated when the first anagram was projected. However, it is possible that the DRN will not deactivate between sequences during the discussion phases with the participant (checking answers, displaying scores). A continuous acquisition of data over time could lead to a better model to be set up in order to correctly predict DRN activation and deactivation during LH experiments.

Finally, the RS phase demonstrated that the connectivity with a region of the DMN (the PCC) differed between the two groups. In the LH group, the DRN is much more connected with the PCC than in the control group. This could suggest that LH affects participants even after the resolution period. Interestingly, DMN changes are also observed in psychological disorders such as attention deficit hyperactivity disorder, obsessive-compulsive disorder, post-traumatic stress disorder and schizophrenia (Sha et *al*., 2019). It would be interesting to understand in a later study with a higher degree of confidence (greater z-threshold and cluster correction) by increasing the number of participants to localize with a greater precision the regions that differ between a non-LH group and the LH group. The LH induced by this very simple experiment shows that the consequences of experiencing unpleasant events ranging from a simple frustrating experience to a true traumatic experience is probably more of a spectrum. Indeed, it’s possible that this simple experience, even if it induces less damage to individuals than a traumatic experience, still induces a slight modification of connectivity in the RS for a short time period. Interestingly, several regions affected by post-traumatic stress disorder (PTSD) such as the vmPFC and caudate nuclei (Fenster et *al*., 2018) are also heavily involved in the controllability detection phenomenon as suggested by Maier et al., 2016 (Maier & Seligman, 2016). Future studies could further increase the time between LH-inducing tasks and the rs-fMRI to determine for how long RS connections differ between the two groups. When a certain period of time between the LH-inducing tasks and rs-fMRI suggests that the LH experiment no longer has any impact on the RS, it would be interesting to repeat the experiment of LH-inducing tasks with the same task several times to see whether increasing the number of LH-inducing tasks can increase the period during which differences are observable on functional connectivity maps in rs-fMRI between control group and LH group. These future experiments could be used to create a model for better predicting and understanding LH generated by events that are non-traumatizing *a priori* but have lasting impacts on individuals.

## 1.6 Conclusion

The findings presented here serve as significant milestones in unveiling the enduring impact of LH on individuals over time. The observed distinctions in resting states between the test and control groups underscore the profound consequences of LH on the structural dynamics of the brain. This study suggests that LH has the potential to disrupt the intricate connections between various brain regions. Consequently, it becomes evident that beyond its initial unpleasant and seemingly virtual nature, LH possesses the capacity to inflict long-term harm on individuals. This crucial insight into the enduring repercussions of LH carries profound implications for the management of individuals within organizational contexts. Acknowledging and addressing the potential neurobiological alterations induced by learned helplessness becomes imperative for fostering a healthier and more resilient workforce.

## 1.7 Acknowledgements

We extend our appreciation to the staff at the Unité de Neuroimagerie Fonctionnelle of the Centre de recherche de l’Institut universitaire de gériatrie de Montréal for their support and collaboration, as well as the Natural Sciences and Engineering Research Council of Canada (NSERC), Discovery Grant number RGPIN-2017-05632. We would also like to thank Jan Valosek, who was a great help in drafting the methodology of the fMRI and rs-fMRI data.

